# Participatory development of innovation and implementation strategy – a practical approach

**DOI:** 10.1101/2025.10.19.679982

**Authors:** Miae Lee, Lesley A. Anderson, Emilia Piltonen, Abha Maheshwari, Julian L. Griffin, Seshadri S. Vasan

## Abstract

**Background:** Healthcare and academic institutions face growing challenges in strategic planning due to rapid advances in medicine and technology, alongside fiscal and workforce constraints that limit traditional consultation. Participatory approaches offer a way to integrate diverse stakeholder perspectives under these constraints, generating contextually relevant strategies that can indicate whether current directions are appropriate or whether priorities have been overlooked.

**Methods:** A structured participatory workshop was conducted at the 10^th^ Grampian Research Conference (June 2025). One hundred seventy-eight participants including National Health Service (NHS) staff, academics, industry partners, patients, and public contributors, engaged in 14 parallel roundtable discussions. Contributions were captured using posters and Post-it notes, collecting 148 written annotations. Data were analysed using thematic and content analysis, supplemented by strategic frameworks including Strengths, Weaknesses, Opportunities and Threats (SWOT/TOWS), and Easy Wins, to identify and prioritise actionable strategies.

**Results:** Five core themes emerged: (1) access to healthcare and services, (2) patient and public involvement and engagement, (3) digital health and service delivery innovation, (4) data access, integration, and governance, and (5) workforce development and culture. SWOT analysis identified strengths in telemedicine, interdisciplinary student training, and patient and public involvement, alongside weaknesses in fragmented data, referral tracking, and workforce pressures. TOWS matrix produced strategy-oriented recommendations such as AI-enabled scheduling, remote monitoring, and transparent referral systems. Easy Wins framework assessment highlighted immediate, low-cost improvements including identifiable NHS caller identification, automated text message reminders, updated informational videos and multilingual materials.

**Conclusion:** By combining participatory outputs with structured strategy tools, this approach demonstrated a resource-efficient model for adaptive planning. The findings align with and extend current national health policy frameworks, offering a replicable approach for institutions aiming to obtain meaningful stakeholder engagement despite fiscal and temporal constraints.

## Background

Healthcare organisations face mounting pressure to adapt strategies in response to rapidly evolving scientific, technological, and societal changes. Advances in genomics, artificial intelligence, and data science are reshaping models of care, while the pace of clinical practice updates increasingly demands rapid integration into everyday care delivery. Traditional five-year strategic planning cycles commonly used in hospitals, universities, and research institutes, struggle to remain relevant and often fail to keep pace with shifting priorities and emerging innovations.^1–3^ At the same time, clinicians, researchers, patients, and other stakeholders operate under significant time constraints and workload pressures, limiting their ability to engage in lengthy or repeated consultation processes.^4,5^ This tension underscores a critical implementation challenge: designing engagement processes that are responsive, efficient, and inclusive while remaining feasible within fiscal and temporal constraints.

Participatory approaches, including co-production and co-design, have emerged as promising methods for developing health interventions and strategies that are contextually relevant and more likely to be adopted in practice.^6,7^ Involving diverse stakeholders, from clinicians and academics to patients and the public, can enhance the legitimacy of decision-making, surface unmet needs, and foster shared ownership of implementation outcomes.^8^ However, participatory processes are resource-intensive, often requiring multiple workshops, facilitated by team-building exercises, or in-depth stakeholder assessments.^9^ In publicly funded systems such as the National Health Service (NHS) Scotland, where resources and workforce capacity are constrained^5^, there is a need for scalable alternatives that retain the benefits of inclusivity while operating within limited time and budget.^10^

The science of teamwork offers insights into maximizing value from constrained engagement opportunities. For example, Belbin’s team role theory emphasizes that groups function most effectively when members adopt complementary roles.^11^ Similarly, the Institute for Healthcare Improvement (IHI) working styles framework provides a streamlined alternative for categorizing group dynamics.^12^ While valuable, these approaches can be difficult to implement in large-scale, real-world settings where participants cannot be pre-screened or purposefully allocated into teams. Thus, there is a practical need for flexible, low-cost methods of organizing short-term collaborative activities that nevertheless generate actionable outputs.

Implementation science offers useful tools for addressing this challenge. Frameworks such as Strengths, Weaknesses, Opportunities and Threats (SWOT) analysis and its strategic extension, the Threats, Opportunities, Weaknesses and Strengths (TOWS) matrix, can be adapted to participatory contexts to translate stakeholder perspectives into actionable strategies.^13^ Similarly, the concept of low cost, high impact “easy wins” resonate with the need to deliver visible improvements while larger scale reforms are underway.^3,10^ Applying these structured but flexible methods to participatory workshops may therefore provide a feasible pathway for generating strategy-oriented insights that are both practical and aligned with health system priorities.

Against this backdrop, we examined whether a structured, time-limited participatory workshop format could generate implementable strategies to guide healthcare innovation in Scotland. Grounded in implementation science principles, we tested whether brief, interdisciplinary discussions supported by systematic capture and analysis of outputs could identify both long term priorities and immediate easy wins (*Figure 1*). We further assessed how these insights align with, extend, or highlight gaps in existing NHS Scotland policies, including *Realistic Medicine*, the *Digital Health and Care Strategy, Scotland’s AI Strategy*, and the *Triple Helix* priorities of NHS-academia-industry collaboration.^14-17^

**Figure 1.**
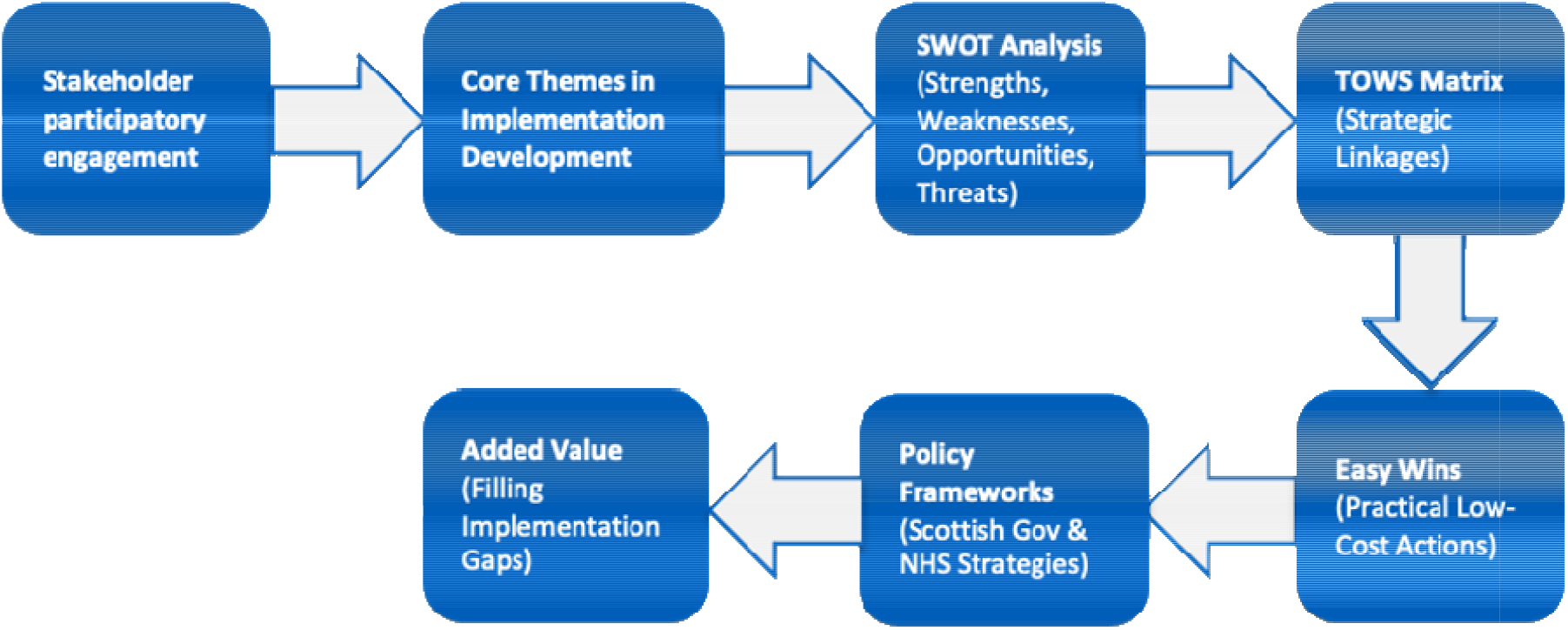
Integrating participatory outputs with strategic planning tools in healthcare implementation

## Methods

### Rationale

This study was situated within the context of the 10^th^ Grampian Research Conference, *“Breaking Traditional Disciplinary Boundaries”*, held on 27-28 June 2025 in Northeast Scotland. The event was designed as a participatory implementation strategy to foster cross-sector dialogue and identify determinants of healthcare innovation adoption. Bringing together NHS Scotland healthcare stakeholders, clinical staff, academic researchers, patients, digital innovators, and members of the public, the conference sought to bring diverse teams involved in healthcare together to explore barriers, facilitators, and strategies for implementing evidence-based and emerging innovations in healthcare delivery.

### Study Design and Participants

The programme included six five-minute plenary presentations highlighting healthcare innovations and lived patient experience, followed by a panel discussion. Presentation topics included drone healthcare technology to improve equity and access, home testing initiatives, the use of artificial intelligence (AI) in cancer screening and pathway redesign, AI-enabled app for reliable health information, and a young patient’s journey navigating NHS services.

A total of 197 participants registered, with 175 participants at the roundtable discussion session. Attendees were seated at 14 tables with 8–9 individuals. NHS participants were identified by a show of hands, and while no tables were NHS-only, one academic-only table was asked to integrate with others to support interdisciplinary dialogue. Based on observation by the convenor (LAA) and conference organiser (SSV), most participants were actively engaged, contributing diverse perspectives across clinical, academic, industry, patient, and public roles.

### Data Collection

Parallel roundtable discussions were structured around three guiding questions aligned with implementation research constructs:

1. *What’s a key challenge or unmet need in healthcare?*
2. *What’s an innovative idea that could address it?*
3. *Who needs to be involved to make it work?*

Participants documented their ideas in real time on Post-it notes or wrote directly onto table posters. At the conclusion of the session, 14 posters containing participant contributions were collected. The following day the posters were publicly displayed to allow cross-table visibility and reflection. Following the event, all handwritten responses were transcribed verbatim into Microsoft® Excel (Version 16.100.4) for analysis. This approach provided a low-cost but reliable mechanism for capturing participant perspectives while maintaining the participatory ethos of the event.

### Data Analysis and Synthesis

We employed a multi-method analytic approach integrating thematic analysis with conventional and summative content analysis.^18,19^ Two research assistants (ML and EP), blinded to participants and not present at the conference, transcribed and reviewed the dataset to ensure familiarity and minimize bias.

Thematic analysis was first conducted to explore participant perspectives in depth.^18^ ML performed an initial review of all posters, followed by an independent review by EP. Post-it notes and directly written annotations were coded line by line and summarized, with labels assigned to capture emerging concepts from each poster. These codes were then grouped into broader categories, from which emergent themes were developed to identify patterns of meaning across the dataset. This iterative process enabled an in-depth interpretation of participants’ ideas and concerns.

To supplement and validate these findings, content analysis was conducted by ML and summative content analysis by EP.^19^ Conventional content analysis ensured themes remained grounded in the raw data, while summative content analysis quantified the relative frequency of key words, concepts, and categories. This approach provided insight into the salience of specific challenges, innovations, and cross-sector interactions across roundtables.

In addition, each poster was analysed using a SWOT framework (ML and EP in tandem) to identify strengths, weaknesses, opportunities, and threats related to healthcare innovation. Individual poster SWOT analyses were then synthesized across all posters to construct a TOWS matrix, highlighting strategic relationships between internal organizational capabilities and external opportunities and challenges.^13^ This enabled generation of strategy-oriented insights relevant to implementation processes.

In line with implementation science principles of phased adoption, “easy wins”, low-cost and high-impact solutions that could be implemented immediately to deliver visible improvements were identified.^10^

### Rigor and Credibility

Analytic rigor was ensured through multiple stages. Coding was conducted independently by two researchers (ML and EP), with oversight from the lead research team (LAA, JG, SSV). Discrepancies in coding and interpretation (e.g., handwriting legibility or category assignment) were resolved through consensus discussions. Content analysis continued until all annotations were categorized and no new categories emerged, ensuring thematic saturation. The combined use of qualitative interpretation and structured strategy tools strengthened both the credibility and translational relevance of findings.

Coding, theme development, and SWOT/TOWS synthesis were reviewed weekly with the lead research team, incorporating both clinical and academic inputs. Consensus was reached at each stage to ensure analytic rigor, credibility, and relevance to implementation science.

### Ethical Considerations

The conference was conducted as a participatory event rather than a formal research study. Nevertheless, principles of ethical engagement were applied: participation in discussions was voluntary, contributions were anonymized prior to transcription and analysis, no identifying information was retained in the dataset. The North of Scotland Research Ethics Committee confirmed that no ethical concerns were identified, and there were no objections to the publication of the study findings.

### Reception and Participation

The roundtable format appeared to be well received. The convenor (LAA) circulated among tables to check in and provide support, encouraging participation and prompting follow-up discussion to help ensure that all voices were represented.

## Results

### Participant Overview

The roundtable format facilitated contributions from a diverse range of stakeholders, including frontline NHS staff, academic researchers, industry representatives, patients, and external partners (*Table 1*). Of the 197 registered participants, 175 attended the roundtable sessions on the first day of conference. Fourteen roundtables, each comprising approximately 8-9 participants, generated 14 posters containing a total of 148 annotations or written comments, which together formed the dataset for analysis.

**Table 1.**
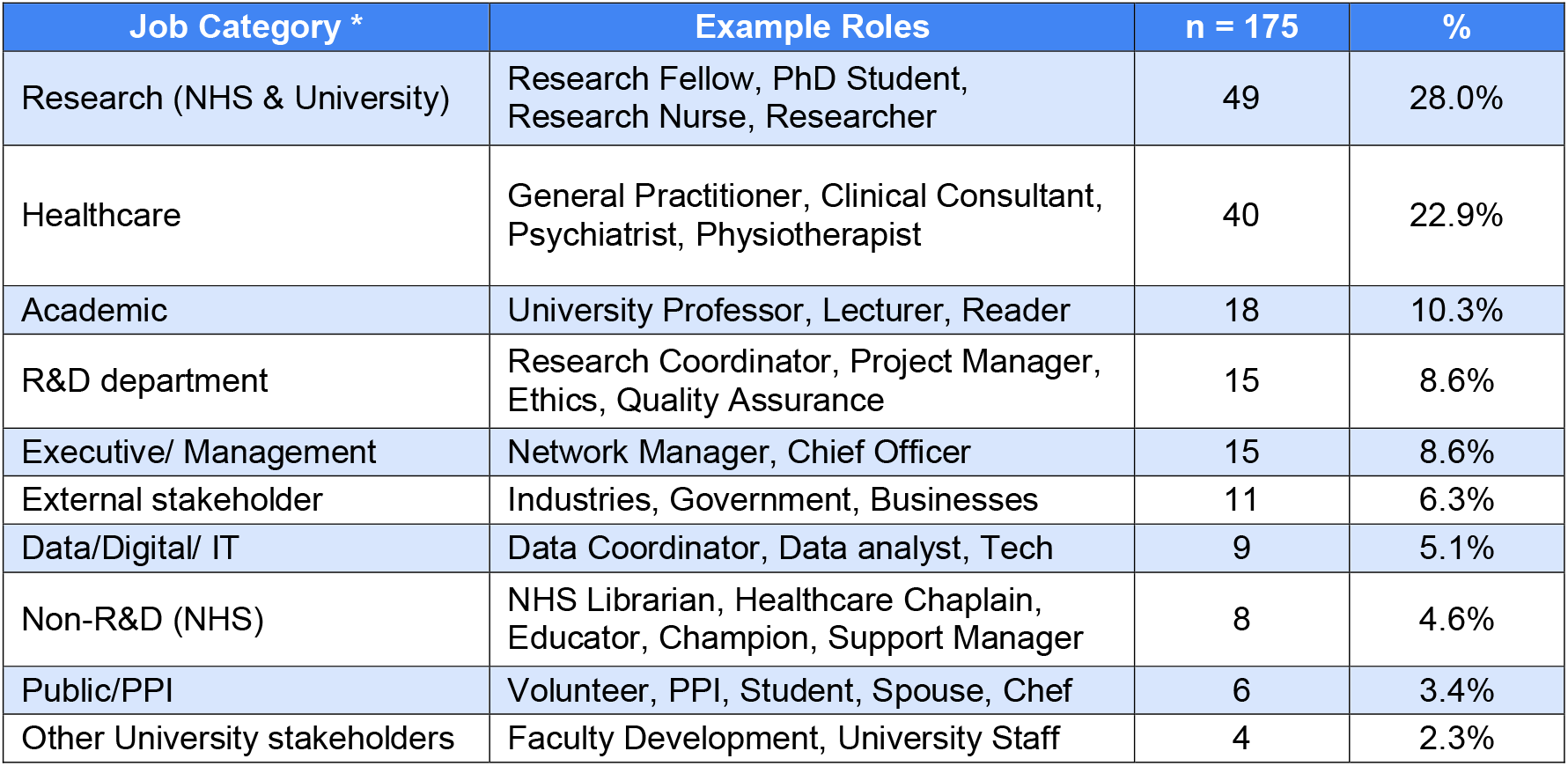
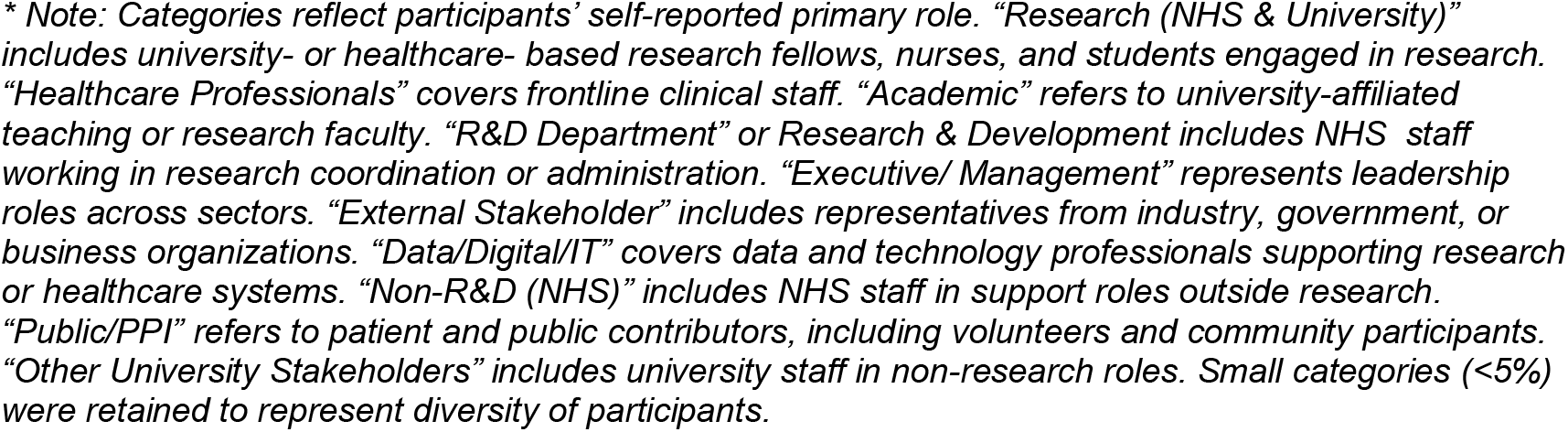
Participant characteristics and stakeholder representation.

### Thematic Analysis: Core Implementation Priorities

Thematic analysis identified five overarching themes, with illustrative examples provided in *Table 2*. Each theme included sub-themes that captured participants’ priorities and recommendations for strengthening NHS service delivery and strategic planning.

**Table 2:**
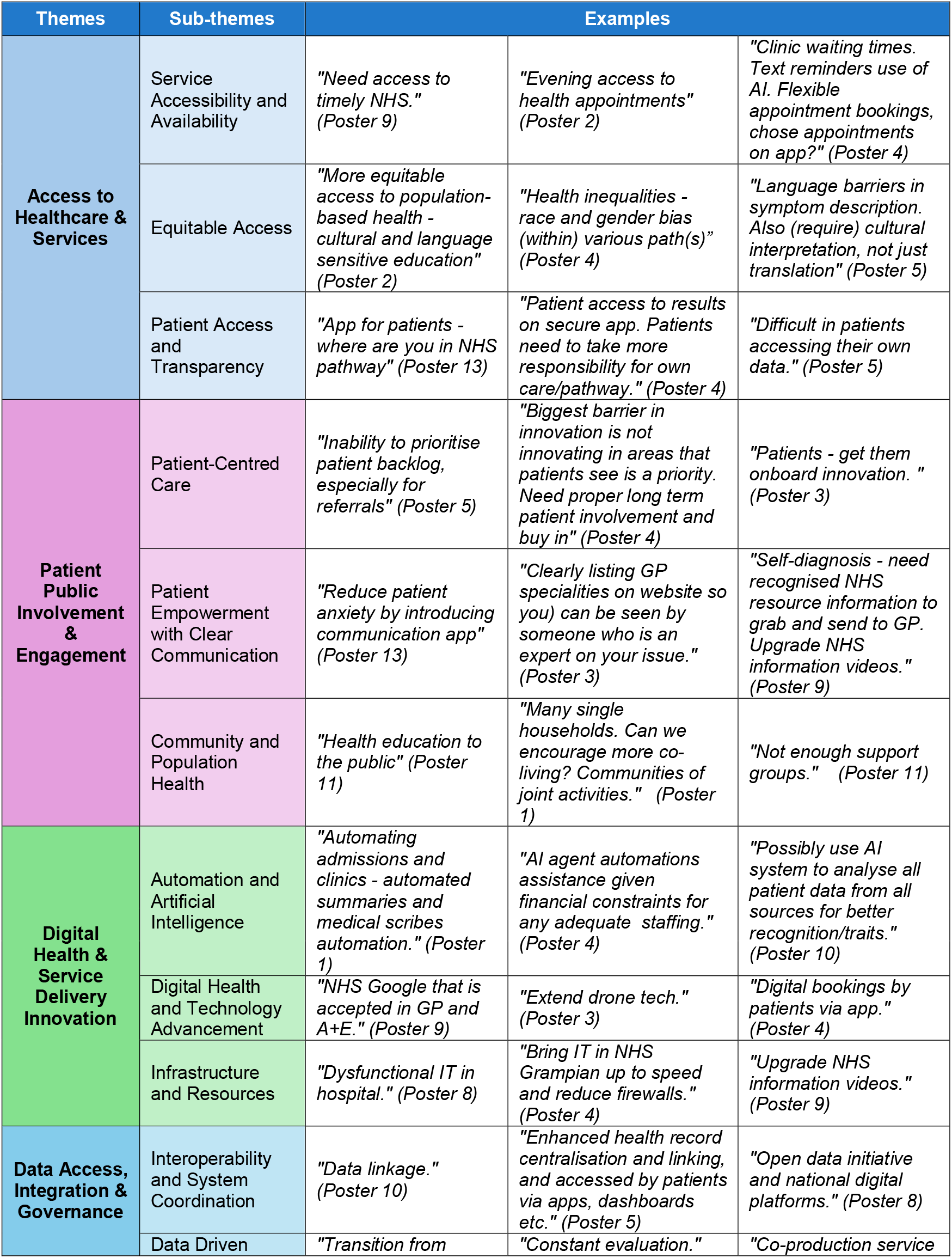
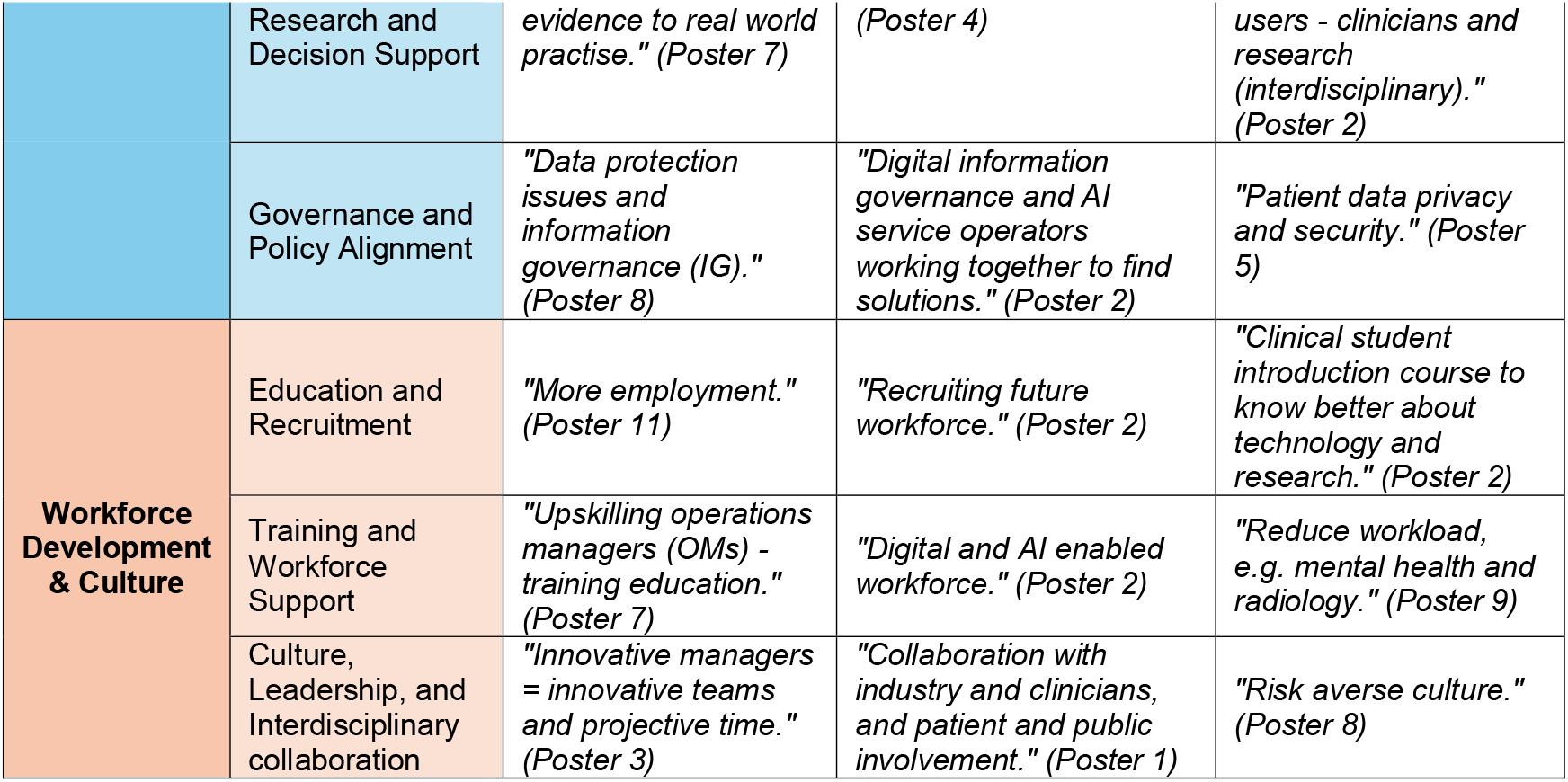
Core themes and subthemes with examples of participant annotations.

#### Theme 1: Access to Healthcare and Services

- *Service Accessibility and Availability*: Participants highlighted the need to enhance timely access to healthcare services through flexible appointments, telemedicine, home testing, visible GP specialties, and efficient care pathways to reduce delays.
- *Equitable Access*: Ensuring fair access was a strong priority, with participants emphasising the need to remove cultural, linguistic and bias-related barriers.
- *Patient Access and Transparency*: Many valued the need to view their health records, referrals, and appointments directly, noting that digital tools could help patients track care in real time and navigate the NHS pathway more efficiently.

#### Theme 2: Patient Public Involvement and Engagement

- *Patient-Centred Care*: Participants emphasized placing patients at the centre of innovation by prioritizing their needs, coordinating care across specialities, centralizing health records, and designing digital systems that simplify navigation care pathways.
- *Patient Empowerment with Clear Communication*: Clear, consistent communication was seen as essential to empower decision-making and self-management, while reducing anxiety. Suggestions included recognizable NHS caller ID numbers, digital reminders, transparent updates on care pathways, and easy tools for appointment management (e.g., cancel, reschedule, or request contact with a healthcare professional).
- *Community and Population Health*: A community-based approach was valued for addressing age-related needs, fostering social innovation, and promoting prevention through education, local support, and coordinated services.

#### Theme 3: Digital Health & Services Delivery Innovation

- *Automation and Artificial Intelligence (AI)*: Participants noted that potential of AI and automation to improve clinical efficiency across admissions, triage, decision support, documentation, scheduling, reminders and wait time management, while emphasizing the need to monitor and address algorithmic bias.
- *Digital Health and Technology Advancement*: Expanding digital access and advancing technological solutions (i.e., drones, digital apps) were seen as opportunities to improve communication, streamline care, and extend healthcare accessibility to a wider community.
- *Infrastructure and Resources:* Modern, reliable IT and physical infrastructure were identified as critical for enabling digital healthcare advancements. Priorities included reducing firewall barriers, investing in system upgrades, and supporting service design that improves data access and hospital operations.

#### Theme 4: Data Access, Integration, & Governance

- *Interoperability and System Coordination*: Participants stressed the importance of standardized, interoperable systems that facilitate communication between services, and enable real-time information sharing, and support open data initiatives for research and service delivery.
- *Data Driven Research and Decision Support*: Strengthening data collection, quality, relevance, and integration across the NHS was seen as key to supporting research, continuous quality improvement, and evidence-informed decision making, with a focus on scalable, cost-effective, high-impact solutions.
- *Governance and Policy Alignment*: Effective innovation was seen as dependent on clear governance frameworks and supportive policies, including automated information governance guidance, open data access, and cross-sector collaboration to ensure responsible technology adoption.

#### Theme 5: Workforce Development & Culture

- *Education and Recruitment:* Participants highlighted the need to prepare the future healthcare workforce by embedding digital literacy, research capacity, and communication skills in early medical education, alongside creating innovative training pathways and strengthening recruitment.
- *Training and Workforce Support*: Upskilling current staff to develop a digitally skilled, AI-enabled workforce was emphasized as a priority. Reducing administrative burden through technology and providing ongoing training and support were viewed as essential for workforce retention.
- *Culture, Leadership, and Collaboration*: A culture of innovation and technology adoption requires engaged leadership and interdisciplinary collaboration across patients, clinicians, industry, and government. Participants noted the importance of overcoming risk aversion, fostering cross-team communication, and embedding innovation into organizational strategies and funding models.

### Summative Content Analysis: Frequency and Salience of Ideas

Summative content analysis identified the most frequently occurring concepts across the dataset, providing insight into both the breadth of issues raised and their relative prominence (*Figure 2*). The most frequently represented theme was Digital Health and Service Delivery Innovation (26.5%), appearing in every roundtable, followed by Access to Healthcare and Services (24.2%), Data Access, Integration, and Governance (20.4%), Workforce Development & Culture (15.6%), and Patient and Public Involvement & Engagement (13.3%). Frequently occurring terms included *data, access, patients, AI*, and *innovation*, reflecting shared priorities across stakeholder groups (*Figure 3*). Less frequent but highly salient terms such as *workforce, culture, infrastructure, and governance* clustered within specific tables, highlighting areas of concentrated stakeholder concern.

**Figure 2.**
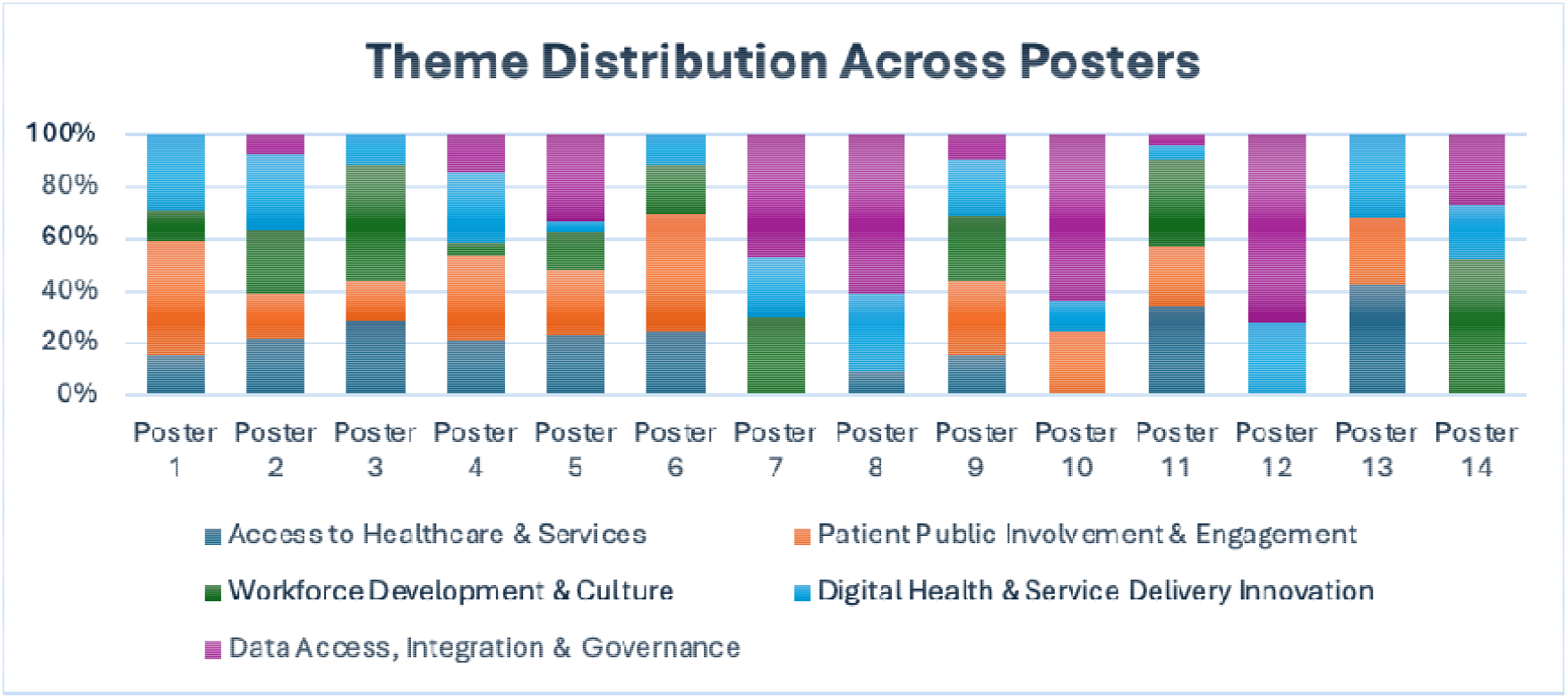
Theme distribution across posters

**Figure 3.**
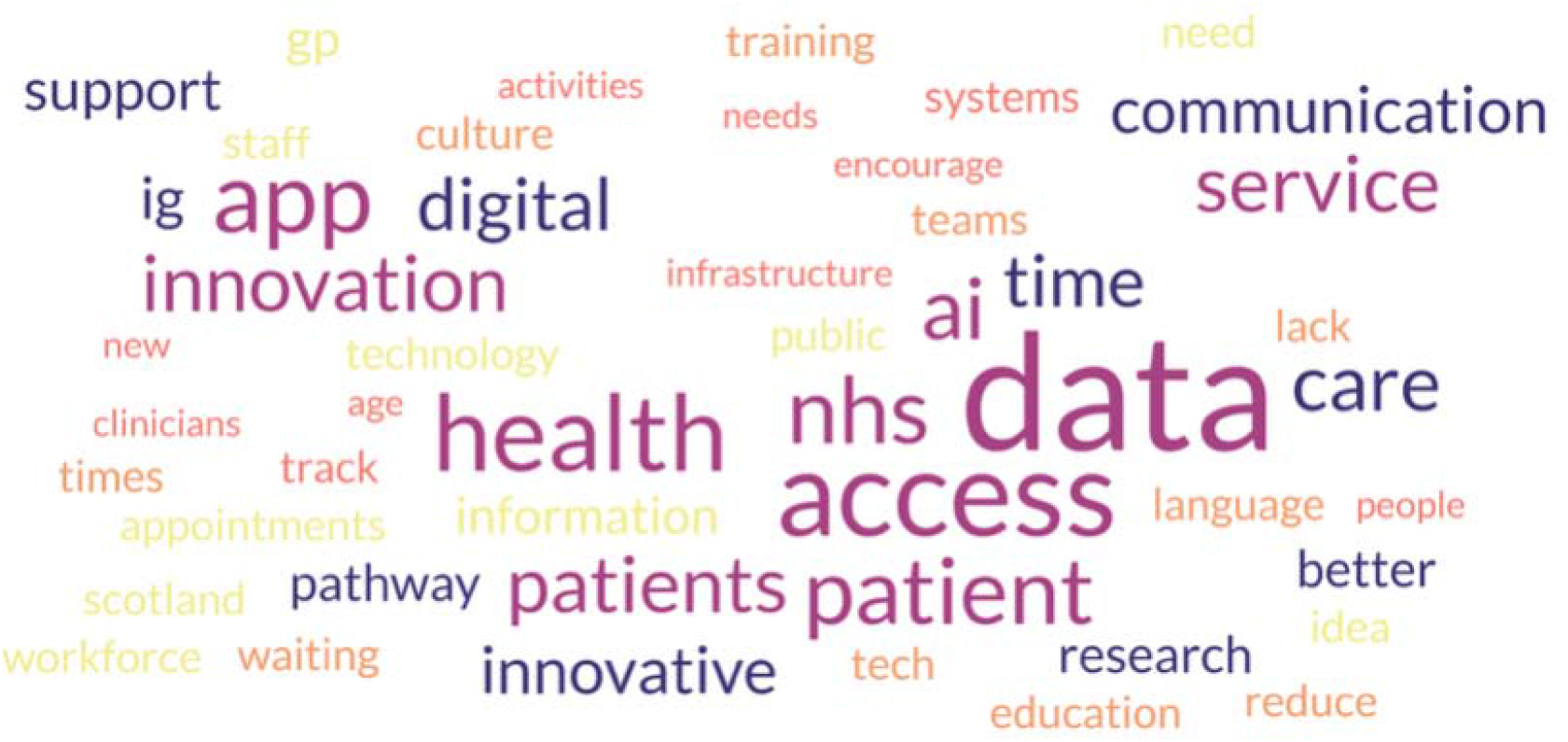
Word cloud highlighting the 50 most frequently used words from the poster quotations

Together, these findings underscore the dual importance of widely shared priorities, such as digital transformation and access, alongside more targeted issues (e.g., workforce culture, infrastructure gaps) raised by specific groups.

### SWOT Analysis: Strategic Landscape of Innovation

Each poster was analysed using a SWOT framework to identify internal strengths and weaknesses and external opportunities and threats, summarized in *Table 3*.

**Table 3:** Stakeholder-derived SWOT analysis summary.

| Internal Strength | Internal Weaknesses |
| --- | --- |
| **Expanded Access to Care:** Telemedicine, e-consults, and NHS care networks, and on-going research in drone delivery extend advanced healthcare to remote patients and smaller hospitals. | **Access and Capacity Challenges:** Delays in care, limited appointment availability, GP/service gaps, workforce shortages, and late-hour access issues |
| **Education and Interdisciplinary Training:** Clinical students gain interdisciplinary training, gaining strong technology and research skills | **Data and Information Gaps:** Fragmented systems, poor interoperability, inconsistent data quality, limited patient access to information, and slow translation of research into practice. |
| **Patient and Public Involvement:** PPI initiatives enable patients and the public to actively shape research and healthcare services, ensuring their perspectives are central to decision-making. | **Communication and Coordination Issues:** Breakdowns between services, teams, and patients; poor referral tracking; and inadequate integration between primary and secondary care |
| External Opportunities | External Threats |
| **Digital & AI-Enabled Transformation:**<br>Leveraging AI, digital platforms, telemedicine, and interoperable systems to streamline care and enhance patient access. | **Resource & Capacity Pressures:** Aging population pressures, staff shortages, burnout, and overloaded services. |
| **Workforce Development & Efficiency:**<br>Upskilling, flexible staffing, innovative job planning, and interdisciplinary collaboration to optimize care delivery. | **Technology & Data Risks:** Slow adoption, infrastructure lags, cyber/privacy risks, digital exclusion, and AI algorithm bias. |
| **Patient Engagement & Data-Driven Care:**<br>Empowering patients through access to records, personalized communication, co-designed services, and data-informed pathways | **Policy, Funding & Cultural Barriers:** Limited funding, governance blocks, policy restrictions, and resistance to change |

- **Internal strengths** included the emerging implementation of new technology in NHS Grampian and the northeast islands, such as telemedicine, e-consults, and integrated care networks, as well as ongoing research in drone-assisted medication delivery. Additional strengths included interdisciplinary education for clinical students and Patient and Public Involvement (PPI). Together, these demonstrate a growing readiness to embed digital tools and patient-centred approaches to service delivery.
- **Internal weaknesses** highlighted persistent system-level barriers, including access and capacity challenges, fragmented data systems, and communication gaps between services and patients. These underscore the need for greater interoperability and service integration to translate innovation into measurable improvements.
- **External opportunities** reflected broader system trends, including digital and AI-enabled transformation, workforce development, and enhanced patient engagement through data-driven care. Seizing these opportunities will require alignment with national policy priorities and sustainable investment strategies.
- **External threats** encompassed resource pressures, technology and data risks, environmental barriers such as air traffic control regulations for drone delivery, and uncertainty in policy or funding landscapes. These external constraints highlight the importance of risk mitigation and equitable implementation planning.

Overall, this analysis provided a structured overview of the strategic landscape for NHS innovation, highlighting both enablers and constraints that must be addressed for effective implementation.

### TOWS Matrix: Translating Findings into Strategy

The TOWS synthesis mapped internal strengths and weaknesses against external opportunities and threats to generate strategy-oriented recommendations, summarized in *Table 4*.

**Table 4:** TOWS matrix linking internal and external factors to proposed strategies.

| TOWS matrix | External Opportunities | External Threats |
| --- | --- | --- |
|  | Digital & AI-Enabled Transformation | Resource & Capacity Constraints |

|  | Workforce Development & Efficiency | Technology & Data Risks |
| --- | --- | --- |
|  | Patient Engagement & Data-Driven Care | Policy, Funding & Cultural Barriers |
| **Internal Strength** |  |  |
| **Expanded Access to Care** | (1) Expand healthcare access with telemedicine, AI-driven triage, and interdisciplinary training for future clinicians<br>(2) Unify care through shared records and interoperability<br>(3) Empower patients with personalized, co-designed, data-driven portals | (1) Deploy telehealth to reduce staff shortage and aging population pressures<br>(2) Leverage tech-skilled students to offset slow adoption risks<br>(3) Strengthen patient engagement systems against policy resistance and digital exclusions |
| **Education and Interdisciplinary Training** |  |  |
| **Patient and Public Involvement** |  |  |
| **Internal Weaknesses** |  |  |
| **Access and Capacity Challenges** | (1) Cut wait time with AI scheduling, remote monitoring, and virtual care<br>(2) Boost capacity with flexible staffing and upskilling<br>(3) Strengthen referrals and coordination through portal tracking and data-driven tools | (1) Prevent bottlenecks with demand forecasting and workforce optimization<br>(2) Mitigate risks through governance interoperability, data quality, and AI validation<br>(3) Advance integrated care pathways with system investment and policy alignment |
| **Data and Information Gaps** |  |  |
| **Communication and Coordination Issues** |  |  |

- **(SO) ‘Maxi-Maxi’ Strategy:** Internal strengths (i.e., existing resources that expand access to care, interdisciplinary training, and PPI) can be harnessed to drive digital and AI-enabled transformation, scaling telemedicine, unifying care through shared records, and empowering patients with personalized, user-friendly portals.
- **(ST) ‘Maxi-Mini’ Strategy:** These strengths also help mitigate resource and capacity pressures by deploying tech-skilled students to accelerate technology adoption and strengthening patient engagement.
- **(WO) ‘Mini-Maxi’ Strategy:** Internal weaknesses (including access and capacity challenges, fragmented data, and communication breakdowns) were addressed through strategies such as exploring AI-driven scheduling, remote monitoring, flexible staffing, upskilling, and transparent referral tracking.
- **(WT) ‘Mini-Mini’ Strategy:** External threats (including resource constraints, technology and data risks, and policy or funding barriers) were countered with recommendations to develop demand forecasting and invest in governance, interoperability, data quality, AI validation, and integrated care pathways.

Overall, the TOWS analysis translated broad insights into strategy-oriented directions for NHS digital, workforce, and patient-centred innovation.

### Easy Wins: Immediate, Low-Cost Implementation Actions

Stakeholders identified several practical, low-cost “easy wins” to accelerate implementation (*Table 5*). Key recommendations included standardizing automated text message reminders across the healthcare system, enabling digital appointment cancellations to reduce no-shows, and providing language-inclusive materials and updated NHS informational videos to promote equity and understanding. Participants also proposed piloting online support groups in high-demand areas such as dementia, chronic care, and mental health, alongside broader public health education campaigns to reduce isolation and ease service pressures. Incremental improvements to patient engagement tools (i.e., adding GP specialty listings, secure medical record access, and referral tracking) were highlighted as ways to empower patients and minimize communication breakdowns. Finally, deploying students in tech-enabled roles, including triage support, digital training, and patient onboarding, was recommended to help alleviate workforce pressures while equipping students with future-ready skills.

**Table 5.** Easy Wins (“Low-Hanging Fruit”) identified by Participants.

| Easy Wins | Description |
| --- | --- |
| **Cutting no-shows** | Roll out automated text reminders, digital appointment cancellation, and NHS recognizable caller ID verification to cut missed appointments and improve trust. These already exists in some clinics and can be standardized across the healthcare system. |
| **Inclusive communication** | Launch language inclusivity measures (translations, plain language materials) and update NHS informational videos for clarity, providing a low-cost, high-impact step toward equity. |
| **Virtual support and education** | Pilot online support groups for dementia, chronic illness, and/or mental health to reduce isolation and ease service demand, combined with public health education campaigns via digital health platforms. |
| **Smarter patient portals** | Enhance existing patient portals, which are not yet available in all regions, by standardizing them across all clinics and simplifying rollout for GP practices, with the goal of national implementation. Over time these portals could expand to include secure access to medical records and a referral tracker outlining each patient's care pathway, helping patients monitor their progress, reduce lost information, and minimize communication breakdowns. Small, incremental improvements should be co-designed with patients to ensure usability and relevance. |
| **Future-ready workforce** | Deploy clinical students with technical skills in tech-enabled roles (triage support, digital training, patient onboarding) to alleviate workforce pressures while building future skills through structured placements or “earn-as-you-learn” pilots. |

Together, thematic analysis, SWOT/TOWS strategy mapping, and easy wins collectively provide a comprehensive, stakeholder-driven actionable framework for strengthening healthcare service delivery through digital innovation, patient-centred care, workforce development, and equitable access.

## Discussion

This evaluation demonstrates that structured, time-limited participatory workshops can efficiently capture strategic priorities and implementable solutions, even within the constraints of limited resources and workforce pressures. By engaging NHS staff, academic researchers, industry representatives, patients, and public contributors, the roundtable process produced a coherent set of implementation strategies that complement and extend national health policies.

Thematic analysis revealed five interconnected themes: (1) access to healthcare and services, (2) patient and public involvement and engagement, (3) digital health and service delivery innovation, (4) data access, integration, and governance, and (5) workforce development and culture. These findings align strongly with existing frameworks and reinforce existing Scottish policies, but also identify critical operational details, particularly around patient-facing digital tools, communication equity measures, and student workforce innovation, that are not yet fully articulated in national strategies. The *Triple Helix* model (link) emphasises cross-sector collaboration: the multi-stakeholder approach adopted here reflects this principle and underscores the value of interdisciplinary co-design in building an innovation ecosystem that can deliver practical, evidence-informed solutions.^17^ The principles of *Realistic Medicine* (link) and the *Population Health Framework* (link) were also evident: participants prioritised transparent care pathways, shared decision-making, and community-based initiatives, and suggested that embedding these values within digital platforms could enhance engagement and patient satisfaction.^14,22^ Scotland’s *AI Strategy* (link) was mirrored in the recognition of AI and automation as enablers of efficiency and decision support, coupled with calls for safeguards to mitigate algorithmic bias and ensure equitable access.^15,16^ These perspectives align with NHS digital governance priorities (link) and the NHS Grampian R&D Strategy (link), which advocates embedding research and innovation into routine practice.^10,20^

The SWOT analysis identified system strengths (e.g., telemedicine, interdisciplinary training, and PPI) alongside persistent system-level weaknesses (e.g., fragmented data, capacity constraints, communication breakdowns). While these broadly align with the transformative aims of Scotland’s *Digital Health and Care* and *AI Strategies*, participants also pinpointed barriers at a more granular level, notably inconsistent referral tracking and the slow translation of research into practice.^15,16^ These issues are not explicitly addressed in national strategies but are beginning to be tackled through initiatives such as the first phase of the new public-facing *Digital Front Door* programme (link), a personalized digital healthcare service, and *Data Safe Haven* platforms (link).^2^ For example Aberdeen’s local *Grampian Data Safe Haven* (DaSH) (link), one of five data safe havens in Scotland, securely links and provides access to pseudo-identified health data for ethically governed research.^21^ Participants also identified PPI, supported through the *Public Involvement Network* (link), which provides regular updates and opportunities for public involvement in research and service design, as a key strength. However, the broader concept of PPIEP (or Patient and Public Involvement, Engagement, and Participation) (link), as defined by the National Institute for Health and Care Research (NIHR) (link), highlights ongoing challenges: while PPI is embedded and required, Patient and Public Engagement (PPE) activities remain limited by budget constraints, with only one event (link) held in the past two years, a challenge shared across wider NHS health boards.^20^ Notably, Participation remains strong, exemplified by initiatives such as Widening Access to Trials in Care Homes (WATCH) project (link), through which NHS Grampian is leading national efforts to involve care homes in vaccine research, strengthen informed consent practices, and contribute to UK-wide guidance in this area.

The TOWS matrix translated these insights into strategy-oriented recommendations. Strengths such as telemedicine and student training pipelines can be leveraged to offset workforce shortages and resource pressures, aligning with the *Triple Helix* principle of joint NHS, academia, and industry investment. Meanwhile, weaknesses in access and data systems could be framed as opportunities for AI-driven scheduling and remote monitoring. These are both being addressed in the *Digital Front Door* programme, supporting the Scottish Government’s *Population Health framework* by promoting equity, efficiency, and accountability.

The identification of “Easy Wins” represents one of this study’s most significant contributions. Stakeholders highlighted low-cost, high-impact actions such as NHS caller ID verification, automated appointment reminders, multilingual patient materials, and updated NHS informational videos. These solutions directly advance the goals of *Realistic Medicine, Population Health Framework*, and the *Digital Health and Care Strategy* (link) by building trust, reducing inequalities, and improving patient experience. Yet they are not explicitly addressed in current policy frameworks, revealing a gap between strategic aims and the “everyday fixes” that matter to patients and frontline staff. Similarly, proposals for structured student involvement in tech-enabled roles illustrate how immediate workforce relief and future digital skills development can be achieved simultaneously, an innovative pathway not fully developed in *Scotland’s Workforce Strategy* (link).

Taken together, the findings show that participatory workshops can serve as a practical method of generating both long-term strategies and immediate operational actions. They bridge the gap between high-level policy and frontline realities, producing outputs that are actionable, resource-conscious, and aligned with implementation science principles of contextual relevance, feasibility, and stakeholder ownership. By highlighting specific operational gaps, patient-facing referral transparency, low-cost communication equity measures, and structured student involvement, this study contributes directly to the implementation literature and offers NHS Scotland actionable levers for accelerating innovation adoption.

### Study Limitations

This study has several limitations that should be considered when interpreting the findings. First, seating at the conference was voluntary. As a result, it was not possible to document which participants sat at which tables or determine whether individuals chose to sit next to friends or colleagues. This limited control over seating arrangements may have introduced bias in the perspectives captured during discussions. Second, feedback collected in participants’ own handwriting proved difficult to interpret at times, raising the possibility of transcription errors or incomplete capture of ideas. Third, the roundtable conversations were not systematically observed or recorded, leaving uncertainty about whether certain individuals dominated the discussions while others contributed less. Finally, innovative healthcare presentations were immediately delivered before the roundtable discussions. While this sequencing may have enriched the discussions by keeping speaker ideas fresh in their minds, it may also have influenced the originality of some contributions.

### Benefits and Reflection for Future

Despite these limitations, the conference successfully fostered strong audience engagement and created opportunities for meaningful dialogue and knowledge-sharing across sectors. Structured opportunities for networking and discussion were incorporated throughout, enabling participants to reflect on system needs and exchange perspectives. Discussions explored policies needed within the NHS, incorporating diverse perspectives from stakeholders across NHS staff, academia, industries, and patients. This collaborative approach aligns closely with the principles of *Realistic Medicine*, particularly its emphasis on shared decision-making, reducing unwarranted variation, and ensuring that care is person-centred and value driven.^14^ Overall, the conference was well-received and offered valuable insights to inform future initiatives.

Reflections on the conference format highlighted several lessons for future events. The use of scribes, rather than reliance on participant handwriting, may improve the accuracy and completeness of recorded feedback. Assigning seating could ensure that participants from diverse roles are distributed across tables, thereby supporting balanced discussions and amplifying diverse voices. In addition, scheduling speaker presentations after round-table sessions may help prevent undue influence on participants, reducing the risk of shaping ideas in advance. Collectively, these adjustments may enhance engagement, inclusivity, and diversity in future conferences.

## Conclusions

Participatory workshops provide a feasible, resource-efficient mechanism for generating implementation strategies in resource-constrained health systems. The combined thematic, content, SWOT/TOWS analyses provided complementary insights into the participatory development of strategies. The innovation conference not only identified key challenges and enablers but also generated actionable strategies tailored to the NHS Northeast Scotland context. By surfacing granular, operational gaps not addressed in current strategies (i.e., patient-facing transparency, equity-focused communication, and structured student involvement), this study demonstrates how participatory approaches can accelerate adoption, foster equity, and improve patient experience. These findings show the value of participatory methods for aligning policy, practice, and patient priorities, and for producing both immediate and longer-term strategies to support sustainable innovation adoption.

## Abbreviations

AI: Artificial Intelligence
NHS: National Health System
SWOT: Strengths, Weaknesses, Opportunities, Threats
TOWS: Threats, Opportunities, Weaknesses, Strengths
ID: Identification
R&D: Research and Development
IT: Information Technology
PPI: Patient and Public Involvement
DaSH: Data Safe Haven
TRE: Trusted Research Environment
PPE: Patient and Public Engagement
UK: United Kingdom

## Acknowledgement

The authors would like to thank Fiona Brebner for her administrative support, Anushree Ganguly and Rituka Richardson for their support with the conference and assistance with de-identified data access and management, and Rachel Hardie for her guidance on ethical considerations. Thanks to conference participants for their contributions to the discussions.

## Authors contributions

LAA, JG, AM, and SSV contributed to conceptualization, methodology, supervision, and project administration (conference organisation, programme development, aims, and research questions). LAA facilitated the roundtable sessions and oversaw data collection. ML and EP conducted data curation and formal analysis, including transcription, coding, thematic analysis, SWOT, and TOWs. ML drafted the manuscript, and EP prepared the figures and tables. All authors contributed to interpretation of the data, critically revised the manuscript, and approved the final version.

